# Negative Electrospray Supercharging Mechanisms of Oligonucleotide Single Strands and G-quadruplexes

**DOI:** 10.1101/2022.07.20.500794

**Authors:** Debasmita Ghosh, Frédéric Rosu, Valérie Gabelica

## Abstract

When sprayed from physiological ionic strength, nucleic acids typically end up with low levels of charging and in compact conformations. Increasing the electrospray negative charging of nucleic acids while preserving the native non-covalent interactions can help distinguish solution folds by ion mobility mass spectrometry. To get fundamental insight into the supercharging mechanisms of nucleic acids in the negative mode, we studied model G-quadruplex structures and single strand controls in 100 mM ammonium acetate. We found that adding 0.4% propylene carbonate, 0.4% sulfolane or 0.1% *m*-NBA induces native supercharging through the charged residue mechanism. However, although 0.4% *m*-NBA shows the highest supercharging ability, it induces unwanted unfolding of solution-folded G-quadruplexes. The supercharging effect resembles the effect of lowering the ionic strength, which could be explained by partial neutralization of the ampholytes when droplets become more concentrated in their non-aqueous components. The supercharging ability ranks: PC < sulfolane < *m*-NBA. *m*-NBA adducts to G-quadruplexes with high charge states confirms that the supercharging agent interacts directly with DNA. Surprisingly, in presence of supercharging agents, more negative charge states also bear more alkali metal ion adducts. This suggests that native supercharging results from larger droplets evaporating to the charged residue, leading to higher concentration of both the supercharging agent and of alkali counterions. However, when negative charge carriers from the electrolyte become too rare, chain ejection accompanied by denaturation, and hence non-native supercharging, can become predominant.

## Introduction

Non-covalent interactions of biomolecules define their structure-function relationships. The goal of “native” electrospray ionization mass spectrometry (ESI-MS) is to preserve weak non-covalent bonds from the solution to the gas phase, by using the least possibly energetic instrumental conditions.^*1*^ However, the charging and desolvation process itself must neither destroy the native non-covalent bonds nor form new ones.

Yet electrospray mechanisms are still debated. The ion evaporation mechanism (IEM) was proposed for low molecular weight species,^*2*^ the charged residue mechanism (CRM) for large globular analytes,^*3*^ and the chain ejection mechanism (CEM) for disordered polymers or unfolded proteins.^*4*^ For proteins, the CEM produces higher charge states than the CRM, and bimodal charge state distributions (CSD) usually indicate that two structural ensembles coexist in solution.^*5*^ For nucleic acids, however, the CSD reflects the adopted electrospray mechanism rather than the solution folding state.^*6*^ For example, nonstructured oligonucleotides adopt only low charge states (following the CRM) at ammonium acetate concentrations of 100 or 150 mM (mimicking the physiological ionic strength), and thus end up compact in the gas phase according to ion mobility measurements.^*6*^

Here, we investigate how supercharging agents (SCA) modify the extent of charging – and possibly the ionization mechanism – of nucleic acids in the negative ion mode. A practical aim is to delineate “native supercharging” conditions, wherein the nucleic acids could gain more charges while still preserving the native non-covalent bonds that were present in solution. A more general aim is to better understand the supercharging mechanisms for nucleic acids in the negative ion mode, and hence progress towards a more generalized understanding of the supercharging phenomenon.

To date, most investigations on ESI charging and supercharging mechanisms were carried out on proteins.^*4,7–10*^ SCAs are less volatile than typical electrospray solvents, and thus concentrate in the evaporating droplets. Thermal or chemical protein unfolding due to droplet heating was suggested early on, but this is incompatible with native supercharging.^*11,12*^ Loo *et al*. argued that the “supercharging” phenomenon must occur in an intermediate regime between the bulk solution and pure gas-phase reactivity, at the charged droplet/air interface.^*7*^ In a recent review, Konermann et al. discussed the mechanisms of protein supercharging in native and denaturing conditions.^*13*^ In native conditions, they attribute supercharging to enhanced charge trapping (decreased evaporation of charge carriers) in the charged residue mechanism as the SCA accumulates upon drying, while in denaturing conditions, the SCA enhances the charging upon chain ejection thanks to their dipole moment. The supercharging mechanism, however, is still unclear for nucleic acids, and for the negative ion mode in general. Xu et al. discussed the supercharging of single-stranded and duplex oligonucleotides in presence of various ionic strength and SCA,^*14*^ but duplex dissociation is observed upon supercharging.^*15*^ An open question is thus whether one can use SCA for native supercharging of nucleic acids.

To test native supercharging of nucleic acids in physiological conditions, we use here G-quadruplex nucleic acid model structures (G4) and unstructured (single stranded) sequence variants of the same length. G4 guanine-rich sequences form secondary structures containing guanine quartets. G-quartets can form from one, two or four strands. The structure is stabilized by the specific coordination of cations in-between stacked G-quartets. The nature and concentration of the cation determines the structure and stability of G-quadruplexes.^16^ Detecting specific cation binding thus indicates that the G-quadruplex was still formed in solution. Stacking interactions, hydrogen bonding, and solvation also contribute to the stabilization of G-quadruplexes. G-quadruplexes are involved in various biological processes such as DNA replication, gene expression, telomere maintenance, and are thus promising therapeutics targets.^17^

To probe ion structure preservation, compaction, or unfolding in the gas phase in supercharging solution conditions, we use ion mobility coupled with mass spectrometry (IM-MS).^*18–20*^ Ion mobility separates ions according to their electrophoretic mobility in a buffer gas (here, helium) under the influence of an electric field. The collision cross section distribution of the ions indicates their gas-phase compactness, and thus allows us to probe the possible conformational changes induced by SCA. We report here that some supercharging solution conditions can lead to native supercharging, and discuss lessons learned on the supercharging mechanisms.

## Materials and Methods

### Sample preparation

All single strands were purchased from Eurogentec (Seraing, Belgium, with RP cartridge–Gold™ purification) and dissolved in water from Biosolve. Desalting is important for our sample preparation since some experiments were done in low ionic strength. We used Amicron ultracel-3 centrifugal filters (Merck Millipore Ltd) with 3-kDa molecular weight cut-off. We first washed the sample several centrifugation passes with 200 mM aqueous NH_4_OAc solution, then several passes with 100 mM NH_4_OAc solution (for ***TG4T***) or water (***20G, 20nonG, 24G, 24nonG***) to reach satisfactory desalting.

All G-quadruplexes (G4) were formed in 100 mM ammonium acetate. The tetramolecular G4 [(dTG_4_T)_4_(NH_4_)_3_] (***TG4T***) was formed from 200 µM single strand, incubated for one week at 4°C. Intramolecular G-quadruplexes d(TTTGGGTGGGTGGGTGGGTT) (***20G***) and d(TT(GGGTTA)_3_GGGA) (***24G***) were formed from 100 µM single strand, incubated for at least 24 hours in the folding buffer (100 mM NH_4_OAc, or 100 mM trimethylammonium acetate (TMAA) with 0.3 mM KOAc). Single strands were annealed at 85°C for 5 minutes to ensure removal of any pre-formed structures. Control experiments were done using non-structured oligonucleotides of the same length: d(TGTGGTGTGTGGTGTGTGGT) (***20nonG***) and d(TGGGATGCGACAGAGAGGACGGGA) (***24nonG***). Supercharging agents were purchased from Sigma Aldrich. They were either 0.4 % (vol) sulfolane (99% purity), m-nitrobenzyl alcohol (*m*-NBA) (≥ 99.5% purity) or propylene carbonate (PC) (99.7% purity), or % *m*-NBA, in 100 mM ionic strength. The DNA structures were injected at 2.5 or 10 μM final concentration for IM-MS analysis.

### UV spectroscopy

Absorbance at 260 nm was recorded using a Uvikon XS and concentrations were determined from the Beer-Lambert law. Molar extinction coefficients were calculated using the nearest-neighbor model and Cantor’s parameters.^21^ The solution stability of the G4 structures was assessed with a UVmc2 double-beam spectrophotometer (SAFAS, Monte Carlo, Monaco), which has a thermostatable 10-cell holder with a high-performance Peltier temperature controller. Experiments were performed by measuring the changes in the absorbance at 295 nm as a function of temperature. Samples were heated to 90 °C. After this, the absorbance was observed at 260, 295, and 350 nm in a series contains a cooling down to 4 °C at a rate of 0.2 °C min^−1^, then heating to 90 °C at the same rate. We converted an absorbance vs. temperature plot into a fraction folded θ vs. temperature.^22^ We fitted upper and lower baseline, where L0_T_ and L1_T_ are baseline values of the unfolded and folded species, then calculate the fraction folded at each temperature, *θ*_*T*_ (Eq. 1). The melting temperature is defined as the temperature at which θ = 0.5.^22^

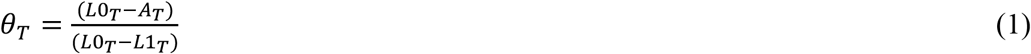

### Electrospray ion mobility mass spectrometry

Structures of intermolecular and intramolecular G4s were probed by IM-MS, performed at 24°C using a 6560 DTIMS-Q-TOF (Agilent Technologies, Santa Clara, CA), with helium in the drift tube. All the experiments were done in negative ion mode with the standard ESI source. Note that the source cleanliness and source settings can significantly influence the allure of the charge state distributions. Unless specified otherwise, the parameters were: fragmentor 320 V, nebulizing gas = 4 psi, drying gas = 1 L/min, trap fill time = 1000 μs, trap release time = 150 μs, and trap entrance grid delta (TEGD) = 2 V. For collision cross section (CCS) verification, [(dTG_4_T)_4_(NH_4_)_3_]^5-^ was injected prior to the analysis, and we checked that we obtained a ^DT^CCS_He_ within 785-791 Å^2^ of its previously determined value (788 Å^2^).^23^ The data were extracted using the IM-MS Browser software version B.08.00 (Agilent Technologies).

The conversion of arrival time distributions to CCS distributions is done as follows.^24^ First, the CCS of the centroid of one of the peaks is determined by the step-field method. Five IM-MS spectra (segments) are recorded with varying drift tube entrance voltages (650, 700, 800, 900, 1000 V), and the arrival time of the centroid of the peak (*t*_A_) is recorded as a function of the voltage difference between the entrance and exit of the drift tube (Δ*V*). A linear regression with Eq. (2) provides CCS from the slope, knowing that (*μ* = *m*_gas_*m*_ion_/(*m*_gas_ + *m*_ion_), *p*_0_ is the standard pressure (760 Torr), *p* is the pressure in the drift tube (3.89 ± 0.01 Torr), *T*_0_ is the standard temperature (273.15 K), *T* is the temperature in the tube (296.15 ± 1 K), *N*_0_ is the buffer gas number density at *T*_0_ and *p*_0_ (2.687 10^25^ m^−3^), *L* is the drift tube length (78.1 ± 0.2 cm), *k*_B_ is the Boltzmann constant, *z* is the absolute value of the ion nominal charge, and *e* is the charge of the electron.

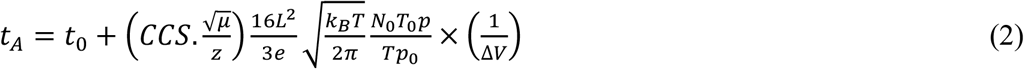

To reconstruct all CCS distributions from the arrival time distributions, the CCS determined with the step field method and *t*_A_ determined for the 650 V segment are used to determine the factor *a* using Equation (3), which is then used to change the axes from *t*_A_ (measured at 650 V) to CCS for all other peaks.

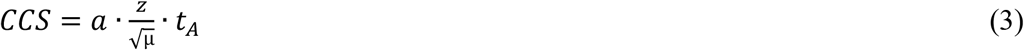

### Calculation of gas-phase structures and collision cross sections (CCS) by molecular dynamics

Molecular dynamics simulations were used to calculate theoretical collision cross sections of the gas-phase structure of the intramolecular G-quadruplex ***20G***. The structure was build using the NMR structure of a propeller-type parallel-stranded G-quadruplex (pdb 2LK7).^25^ One thymine on the 5’ and 3’ end was added. To reach the chosen experimental charge states (5^-^ and 10^−^), protons were added to the phosphate groups). The structures were optimized at the semi-empirical level (PM7) ^26^ using Gaussian 16 rev. C.01.^27^ Then, Atom-Centered Density Matrix Propagation molecular dynamics ^28,29^ (ADMP, 10000 fs, 296 K) at the semi-empirical level (PM7) were performed. The theoretical CCS values were calculated for a structure every 20 fs, using the trajectory model (Mobcal, original parameters for helium).^30^

## Results and Discussion

### G-quadruplex solution structures can be natively supercharged

We studied here several G-quadruplex nucleic acid structures: the tetramolecular G-quadruplex ***TG4T*** with 3 NH_4_^+^, the parallel-stranded intramolecular G-quadruplex ***20G*** with 2 NH_4_^+^, and the hybrid-folded ***24G*** with 2 NH_4_^+^. ***20G*** is 99.99% folded and ***24G*** is 98.48% folded in 100 mM aqueous NH_4_OAc at 22°C (the melting curves are shown in Figures S1-S2). We also used single strand counterparts that cannot fold into G-quadruplexes because they lack enough contiguous G-tracts: ***20nonG*** and ***24nonG***, with no cations specifically bound.

Figure 1 (top row) shows spectra obtained for the G-quadruplexes ***TG4T, 20G*** and ***20nonG*** in 100 mM aqueous NH_4_OAc. In high ionic strength without SCA, structured and non-structured oligonucleotides have a similar CSD: the most abundant charge states for sequences containing 20 or 24 nucleobases are 4- and 5-. The charging occurs according to the charged residue mechanism (CRM), irrespective of the solution folding state.^6,14^

**Figure 1.**
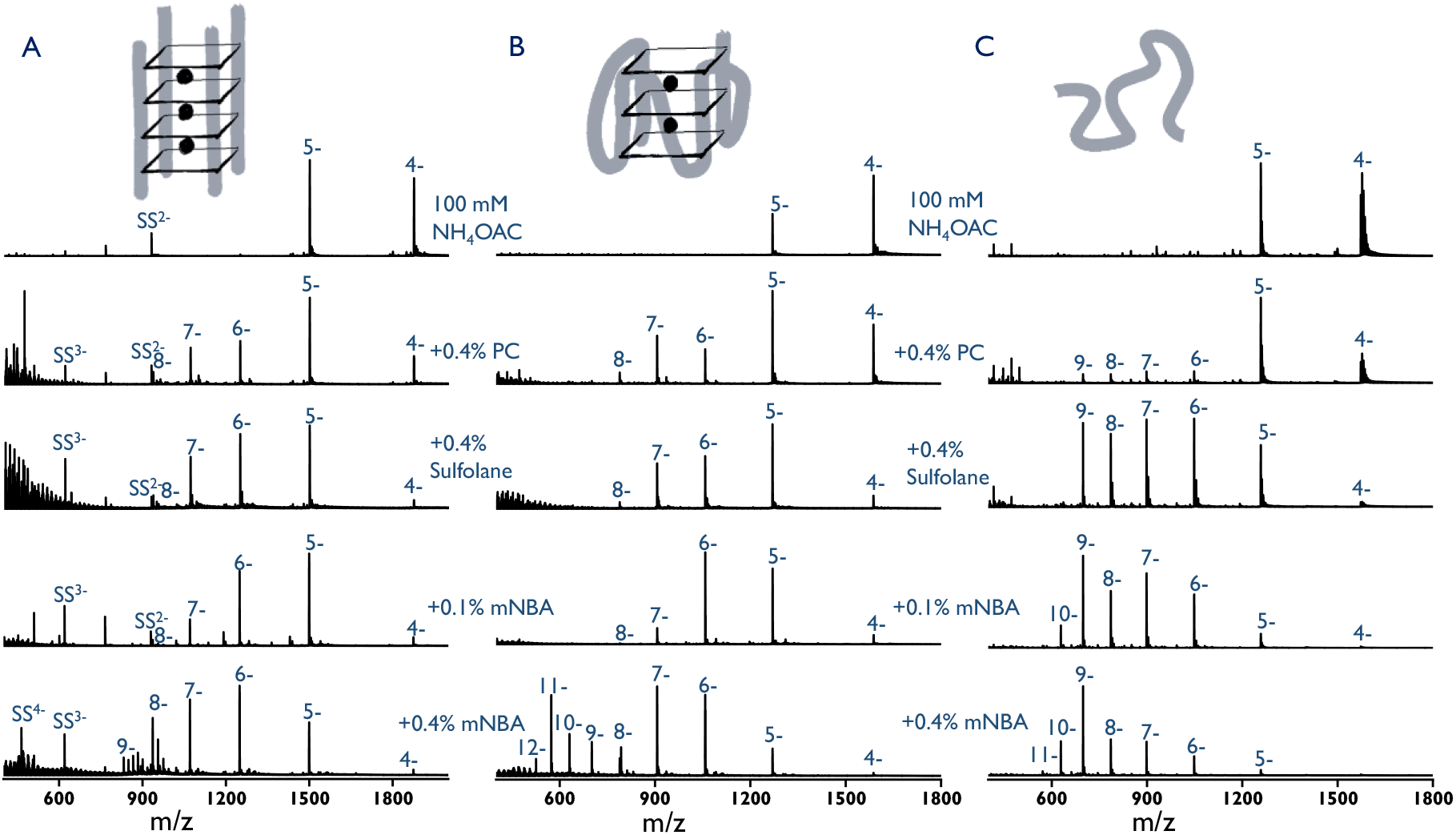
ESI-MS of oligonucleotide structures (10 µM each) in 100 mM aqueous NH_4_OAc, without the addition of SCA (top row) or with 0.4% PC, 0.4% sulfolane, 0.1% and 0.4% and *m*-NBA. (A) Intermolecular ***TG4T***; (B) intramolecular ***20G***; (C) non-structured ***20nonG***.

We observe higher charge states when SCAs such as PC, sulfolane or *m*-NBA are added. For 0.4% of PC, 0.4% sulfolane or 0.1% of *m*-NBA, we obtain charge states up to 8- for ***20G*** and 9- for ***20nonG***. In 0.4% *m*-NBA, the CSD depends on the structure bottom row of Figure 1): ***TG4T*** (the most stable G4) keeps a unimodal CSD, the folded ***20G*** shows a bimodal CSD (with a second, high-charge distribution centered on 11–), while the nonstructured control ***20nonG*** has mainly a monomodal distribution centered on high charge states.

Chen and coworkers observed that, at equal amounts, sulfolane induces less charging than *m*-NBA, but the effect of the *m*-NBA amount was not studied.^14^ With proteins, different amounts were used in different studies, and the results depend on the molecular system. For example, using increasing amounts of *m*-NBA disruption of holomyoglobin into apomyoglobin starts at 0.4%,^11,12,31^ concanavalin A dimers in 200 mM NH_4_OAc are disrupted at 1% *m*-NBA,^32^ whereas several systems are undisrupted even at 1% m-NBA.^33^ Note that compared to sulfolane and PC, the solubility of *m*-NBA in water is low, around 0.05 % in volume.^34^ Although the solubility may differ in presence of electrolyte, solubility might be at the origin of the difference observed with *m*-NBA compared to sulfolane or PC.

Here we distinguish three ranges of charge states: (i) the initial “low” charge states in absence of supercharging agents (4– and 5–), (ii) a “medium” charge state distribution centered on 6– and 7–, and (iii) a “high” charge state distribution, centered on 9– for ***20nonG*** and 11– for ***20G***. PC and sulfolane mostly promote the second distribution, and *m*-NBA is best at promoting the third distribution. Also, the amount of the third distribution depends on the solution structure: it is fully promoted only for the unstructured oligonucleotides, in which case we suspect a production of these charge states via a chain ejection mechanism.

To characterize which charge states retain a native structure in the gas phase and find whether supercharging conditions are native, we use two observables: the retention of specific NH_4_^+^ ions in the complexes, and the gas-phase shape (compact or extended) inferred from ion mobility measurements. Figure 2 compares the collision cross-section distribution (CCSD) of the mass spectra presented in Figure 1. As expected, based on Coulomb repulsion, the average CCS increases with the number of charges, but groups also emerge. Charge states 4– and 5– obtained without SCA are always compact (below 800 Å^2^). A preserved quadruplex with thymine overhangs folded back in a compact manner has a calculated CCS_He_ of 780 Å^2^ (from in-vacuo MD at the PM7 level and TM for CCS calculation, Figure 2D). The histogram of the calculated CCS upon molecular dynamics at the PM7 level is matching extremely well with the experimental distribution for ***20G***^5–^ (Figure S3).

**Figure 2.**
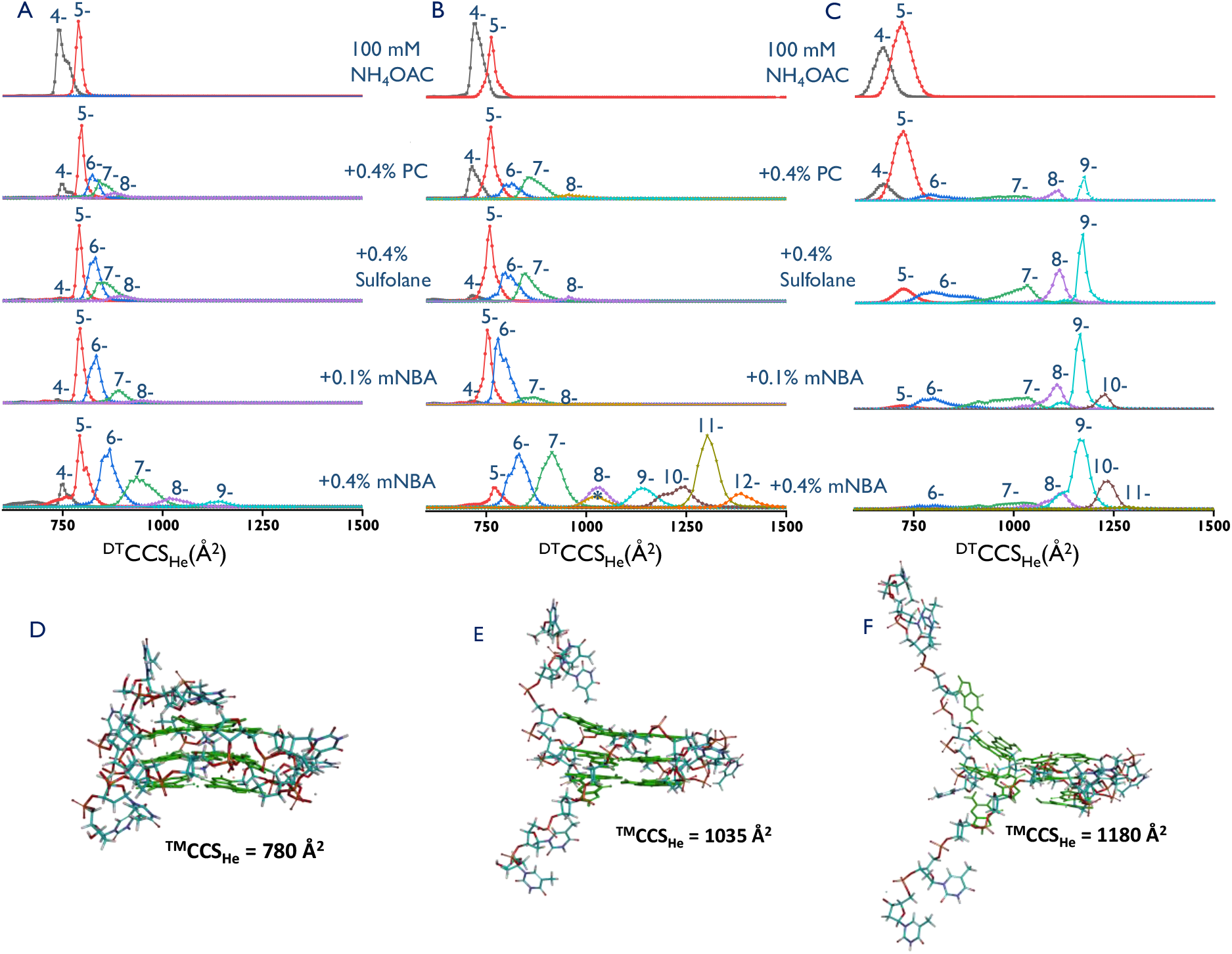
CCSD for ions of oligonucleotides in 100 mM NH_4_OAc, without the addition of SCA (top row), with SCA, 0.4% PC, 0.4% sulfolane, 0.1% and 0.4% and *m*-NBA. A-C shows the difference in ion conformations induced by charging and supercharging of intermolecular ***TG4T***, intramolecular ***20G***, and non-structured ***20nonG***, respectively. For ***20G***, 4– to 7– conserved 2 specific NH_4_^+^, 8-represent with and without 2 specific NH_4_^+^ (*), and 9– to 12– ions contain 0 NH_4_^+^. D-F shows calculated structures in-vacuo (PM7 and TM for CCS calculation). Structures for 5– with 2 inner NH_4_^+^ (D), two shapshots of the 10– simulation without inner NH_4_^+^ (E, F).

For charge states up to 7–, the average CCS remains below 900 Å^2^ for structures that are folded in solution. In *m*-NBA, the highest charge states with preserved inner ammonium ions (preserved G-quadruplex) is 8–, and molecular modeling shows that if G-quartets are preserved but thymine overhangs are extended, the CCS is 1035 Å^2^ (Figure 2E). This suggests that for charge states 4- to 8- (the first two distributions, defined above as “low” and “medium” charge states), native G-quadruplex conformations are preserved from solution to the gas phase. Thus 0.4% PC, 0.4% sulfolane or 0.1% *m*-NBA induce native supercharging from solutions at physiological ionic strength.

In contrast with ***20G***, the charge states 7– to 9– produced from the unstructured ***20nonG*** have higher CCS. This may be due either to formation via the CEM or to formation via the CRM followed by gas-phase unfolding, because the Coulomb repulsion is not counteracted by a strongly specific network of native bonds. In these conditions, although the CSDs are similar for intramolecular G-quadruplexes and unstructured single strands, the CCSD produced by these medium charge states allows us to differentiate between the folded and unfolded solution structures.

The situation is different in 0.4% of *m*-NBA. The CCS distributions are broader and displaced towards larger CCS (see charge states 6– or 7–, the possible reasons will be discussed further below). The ***TG4T*** quadruplex can hardly be charged beyond 8–, and the average CCS_He_ thus remains below 1000 Å^2^. The three inner NH_4_^+^ cations are preserved. Thus, in that case, native supercharging (CRM) may still be at stake. In contrast, the nonstructured oligonucleotide produces almost exclusively high charge states and conformations larger than 1100 Å^2^, implying a disruption of the G-quadruplex in the gas phase (see the model at 1180 Å^2^ generated for ***20G***^10–^, Figure 2F). If denaturation has occurred before the acquisition of the final charge state, the ionization must have followed a chain ejection mechanism. We will come back to this point in a subsequent section.

The shift or not of charge state distributions and CCS distributions in 0.4% *m*-NBA appears as potentially a good way to assess solution structures. However, the results for ***20G*** in *m*-NBA warn against any quantitative interpretation: although the fraction folded in solution is higher than 99.99% (Figure S1), ***20G*** shows a bimodal CCSD, with a significant fraction of ions following the CEM. Also, starting from charge state 8–, inner NH_4_^+^ cations are lost (Figure S4), indicating the disruption of the G-quartet core. Similar results were obtained for ***24G*** and ***24nonG*** (Figure S5 and S6). Thus, using 0.4% *m*-NBA can result in non-native supercharging as well as native supercharging.

### Supercharging with additives vs. by lowering the ionic strength

Previous works described the effect of ionic strength on the electrospray charging process,^6,35^ and Xu *et al*. suggested that the effect of adding supercharging agents on duplexes was analogous to the effect of lowering the ionic strength.^14^ To explore this on our systems, we recorded ion mobility mass spectra at 0.5 to 100 mM NH_4_OAc. Figures 3 and 4 show the results for ***TG4T, 20G*** and ***20nonG***. For intramolecular 20-mers (containing 19 phosphate groups), the CSD changes in three steps: at 100 or 50 mM the CSD has only low charges (4– and 5–), between 10 and 1 mM a second set of medium charge states (from 6– to 9–) appears and around 1 mM a third high-charge distribution (centered on 11– and 12–) takes over. Figure S4 shows that charge states up to 7– retain their inner ammonium ions. The ***TG4T*** quadruplex dissociates in solution below 1 mM NH_4_OAc. Non-structured oligonucleotides showed more clearly bimodal distributions between 10 and 1 mM, centered on 5– and 9–, but with few medium charge states. Even higher charge states (up to 12–) predominate around 1 mM (charge states 4– and 5– almost disappear). Thus the “medium” charge state distribution is more prominent when there is some solution structure to be preserved.

**Figure 3.**
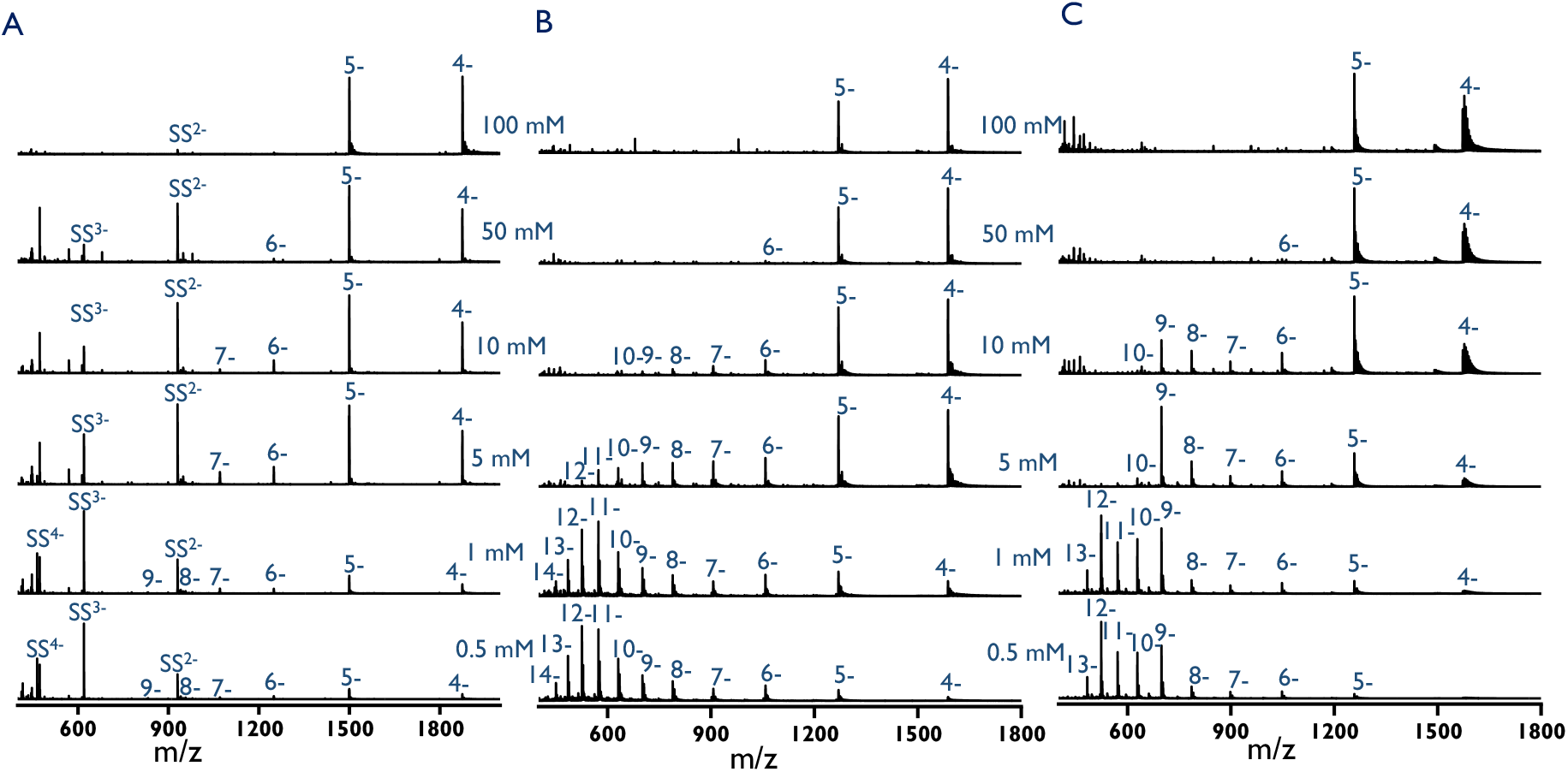
Mass spectra acquired for folded oligonucleotides ***TG4T*** (A), ***20G*** (B), and non-folded ***20nonG*** (C) in different ionic strengths (using aqueous NH_4_OAc solution). This shows the change in CSD as a function of ionic strengths. Appearance of higher charge states were observed at low ionic strengths.

**Figure 4.**
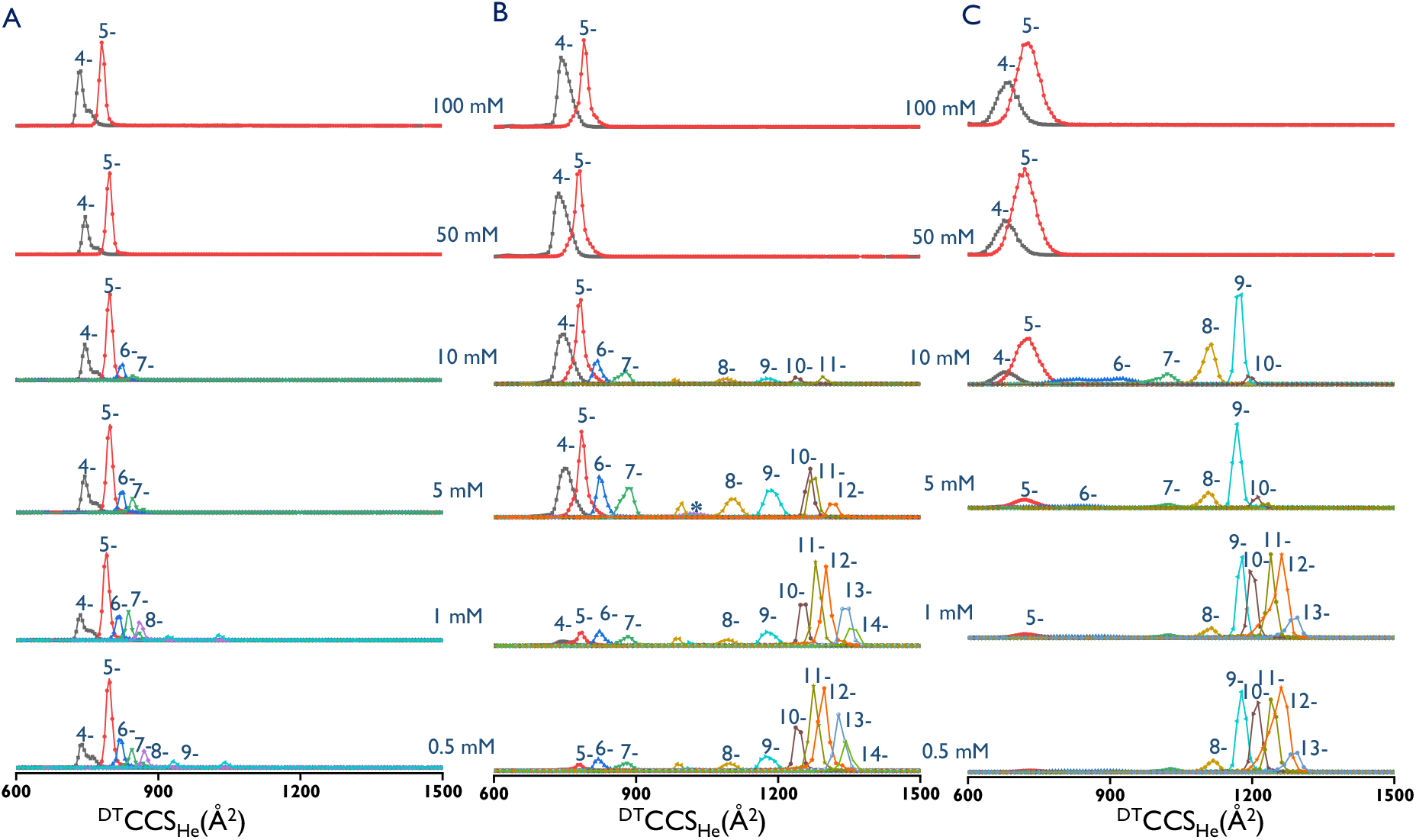
CCSD of folded oligonucleotides ***TG4T*** (A), ***20G*** (B), and non-folded ***20nonG*** (C) in different ionic strengths (using aqueous NH_4_OAc solution). For ***20G***, 4- to 6- conserved 2 specific NH_4_^+^, 7-represent with and without 2 specific NH_4_^+^ (*), and 8- to 14- ions contain 0 NH_4_^+^. This shows unimodal distribution for ***TG4T*** but bimodal distribution for ***20G*** at low ionic strength. 20nonG predominately produce high charge state ions at low ionic strength.

In some respects, adding supercharging agents produces a similar effect as lowering the ionic strength. We thus speculate that SCA clustering around to the analyte could displace the counterions and thus promote high charging (and solution unfolding in extreme cases), but a transition to another charging mechanism (CRM to CEM) also happens at low enough ionic strength, and this is even more prevalent for unstructured strands. In a previous paper,^6^ we theorized that the transition from CRM to CEM for nucleic acids in the negative mode was caused by not having enough negative charge carriers coming from the buffer to the droplet surface, forcing the nucleic acid polyanions to come to the surface, where they can take a chain ejection pathway. We thus studied in more detail SCA and counter-ion interactions with our nucleic acid.

### m-NBA can interact with G-quadruplex nucleic acids

In Figure 1, adducts are visible for charge states 8- and 9- of ***TG4T*** in 0.4% *m*-NBA, and the number and abundance of adducts increases with the charge state. Also, in *m*-NBA, the CCS distributions are broadened and shifted to higher values, and this effect is stronger at higher charge states (Figure 2). We therefore hypothesize that ion mobility peak broadening could be due to *m*-NBA adducts on the DNA persisting in the ion mobility tube, and being partially declustered after the drift tube. To test this hypothesis, IM-MS experiments were repeated with softer tunning parameters in the post-IMS region (the parameters are listed in supporting Table S1, and full discussion of the tuning parameters can be found in reference ^36^). Figures 5 and S6 confirm that even more adducts are detected when using soft post-IMS conditions. This confirms that *m*-NBA interacts directly with ***TG4T. 20G*** showed few adducts with *m*-NBA. We have not seen adducts of sulfolane or PC with oligonucleotides in these conditions.

**Figure 5.**
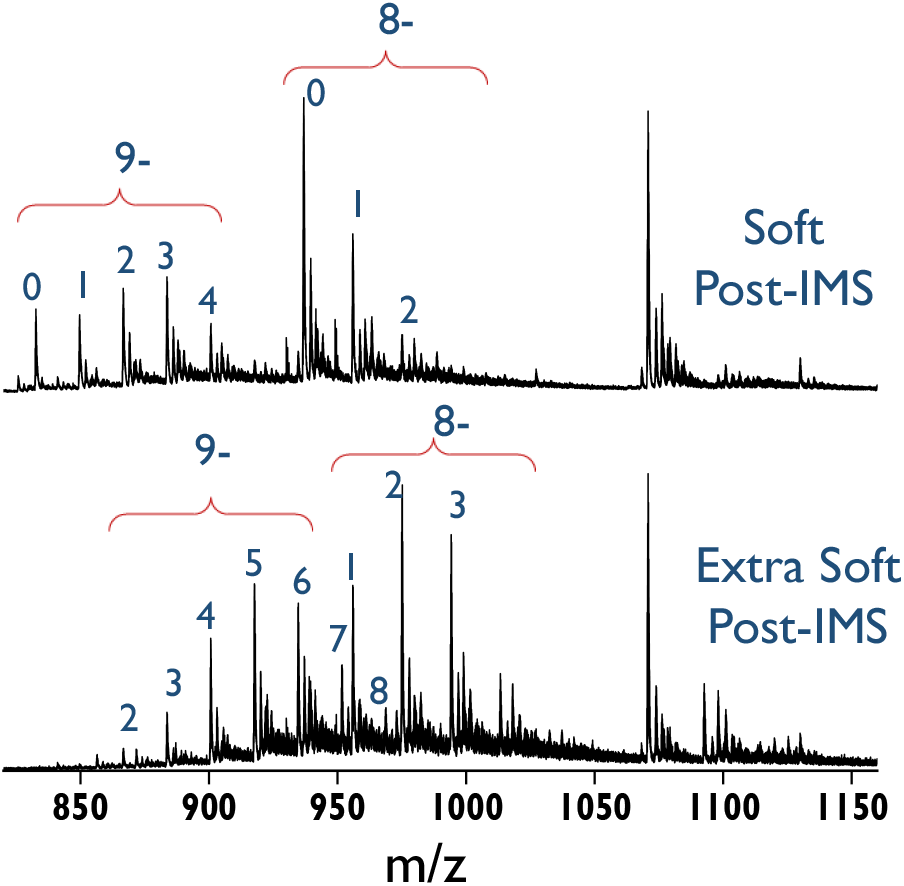
Formation of *m*-NBA adducts with ***TG4T*** in soft post IMS conditions. Zoom on the 9- to 7- charge states. Experiments were done using 0.4% of *m*-NBA in 100 mM aqueous NH_4_OAc. The changes made in the tuning for post-IMS are listed in Table S1.

### Supercharging increases potassium ion condensation on the nucleic acid polyanions

To probe counterion condensation on nucleic acids upon electrospray in supercharging conditions, we need a non-volatile cation. Here we used K^+^ (0.3 mM KOAc), with the 100 mM ionic strength ensured by trimethylammonium acetate (TMAA). G-quadruplexes can form in those conditions, with K^+^ ions in-between the G-quartets,^37^ but extra non-specifically bound cations formed before the acquisition of the final charge state (either in solution or upon charging) are not removed by further collisional activation. Without KOAc, the extent of supercharging is similar in NH_4_OAc and in TMAA (Figure S8).

Figure 6 shows the charge state distributions and zooms on some adducts distributions for ***20G*** (Fig. 6A) and ***20nonG*** (Fig. 6B) in TMAA/KOAc. These results were filtered in IM-MS browser to eliminate single and doubly charged background (Figure S9). For ***20G***, the first peak of the mass distribution indicates that 2 K^+^ ions are specifically bound, even for the highest charge states (9– to 11–) for which the CCS values indicate gas-phase denaturation (Figure S10). This suggests that the K^+^-bound G-quadruplex structure is preserved up to when the final charge state was acquired, and that extension due to Coulomb repulsion occurs after charging. However, given that parallel G-quadruplexes are favored in dehydration conditions,^38,39^ this gives no clue as to how many water molecules were still around at the moment of final charge state acquisition. The non-native high charge states formed in *m*-NBA might be formed from droplets that are very rich in *m*-NBA and contains very few water molecules, yet are capable to take DNA structures along.

**Figure 6.**
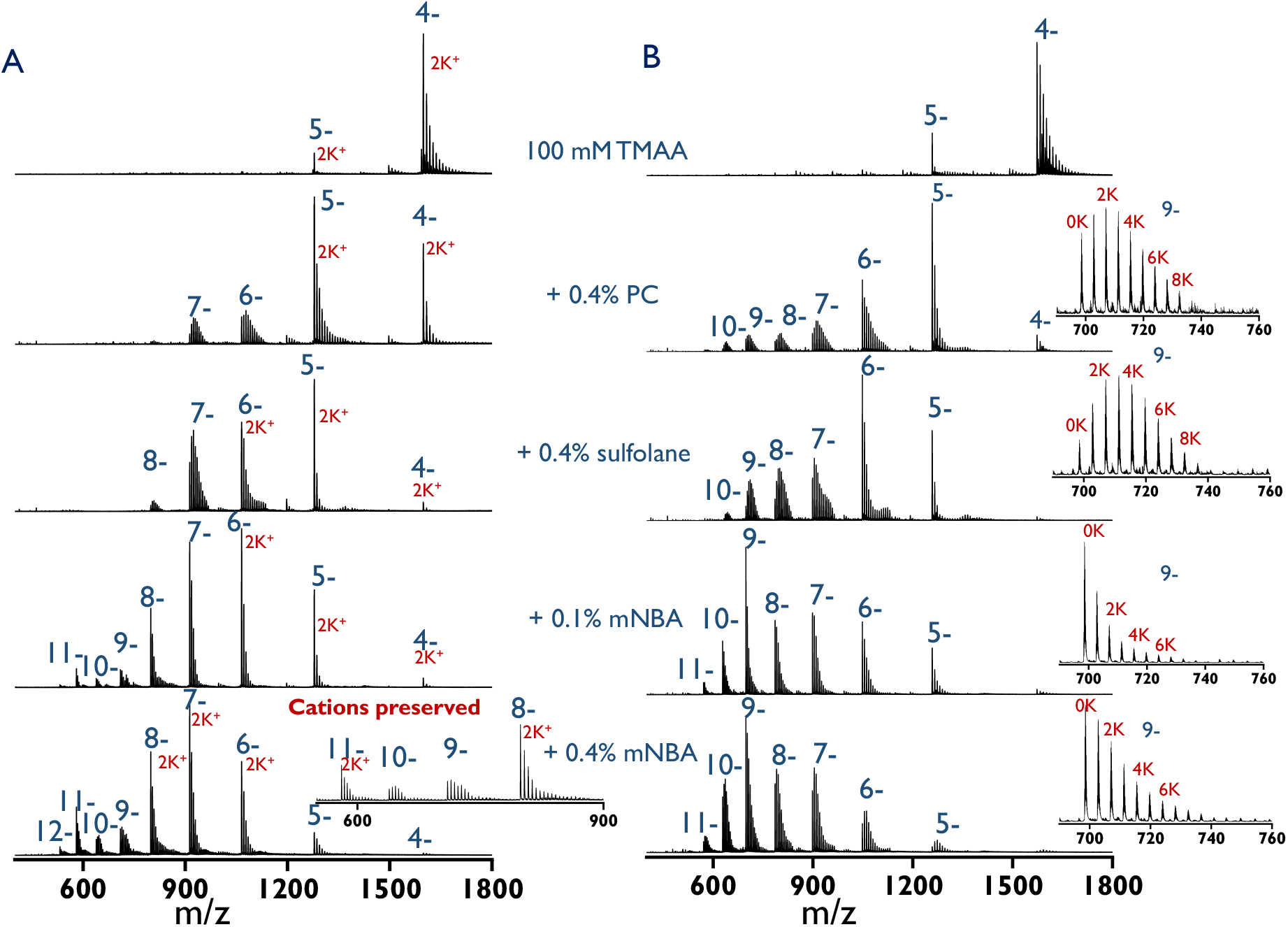
ESI-MS spectra for (A) ***20G*** and (B) ***20nonG*** recorded in 100 mM TMAA with 0.3 mM KOAc, without SCA (top row), and with SCA, 0.4% PC, 0.4% sulfolane, and 0.1% and 0.4% *m*-NBA. The inset in panel (A) shows the preservation of 2 specific K^+^ ions in highest observable charge states in 0.4% *m*-NBA. The insets on panel (B) show counterion condensation for the 9- ions with the different SCAs.

The other intriguing aspect is the potassium adducts distribution, which is better discussed on Figure 6B with the unstructured oligonucleotide ***20nonG***. In negative mode, high charge states should logically bear fewer adducts with alkali metals, as observed without SCA (top row). But this isn’t the case with supercharging agents. In 0.4% PC or sulfolane, K^+^ adducts are more abundant and numerous for higher charge states (i.e., the “medium” distribution). We also note for sulfolane that some K^+^ adduct distributions are bimodal. For *m*-NBA, which produces more of the third, high-charge state distribution, the K^+^ nonspecific adducts are less numerous than in PC or sulfolane and there is not such a significant charge state dependence, but adducts are still abundant. We obtain a similar trend when using NH_4_OAc instead of TMAA (Figure S11).

These observations differ from those reported for native supercharging of positive ions, where both sulfolane and *m*-NBA suppress Na^+^ adducts.^40^ For negative ions, formation of *higher* charge states with *more* K^+^ adducts means either that the supercharging agent displaces TMAH^+^ or NH_4_^+^ from the nucleic acid better than they displace K^+^, or that the acidic form of the ampholyte (TMAH^+^ or NH_4_^+^) is neutralized by the supercharging agents. Furthermore, the large number of nonspecific K^+^ adducts on high charge states indicate that they are issued from large droplets evaporating to dryness, hence suggesting a formation via the charged residue mechanism. When the CEM is made favorable by lowering the ionic strength (Figure S12), the trend in adduct formation becomes again normal, with fewer adducts on high charge states. All evidence points to a different mechanism for native supercharging (CRM) and non-native supercharging (CEM), and that *m*-NBA can induce both native and non-native supercharging.

## Conclusions

Native supercharging of structured oligonucleotides can be achieved by adding 0.4% propylene carbonate, 0.4% sulfolane or 0.1% *m*-NBA to solutions physiological ionic strength. In those conditions, all ions are formed via the CRM, but adding a few more charges can induce expansion upon Coulomb unfolding, which will happen at lower internal energies for ions having no special hydrogen bonding or cation binding network in solution. Native supercharging does not allow to distinguish solution structures based on the charge state distribution alone, but is useful when combined with ion mobility measurements. 0.4% *m*-NBA (which is above the solubility level of *m*-NBA in water) has an even higher supercharging ability, but can cause structures to unfold, and it is still unclear whether this is due to ionization via the charged residue mechanism followed by Coulomb unfolding, or ionization via a chain ejection mechanism. In those conditions, the charge state distributions depend on the solution folding state. The caveat is that 0.4% *m*-NBA causes unwanted unfolding of solution-folded G-quadruplex structures, and quantification of the fraction folded based on deconvoluting the charge state distribution is unreliable.

To some extent, the supercharging effect resembles the effect of lowering the ionic strength, and the supercharging ability ranks: PC < sulfolane < *m*-NBA. Moreover, *m*-NBA can interact directly with G-quadruplexes, forming long-lived adducts that cause a broadening of the CCS distributions. The experiments with non-volatile K^+^ ions show no cation displacement from the DNA by the supercharging agent. In line with a mechanism suggested by Loo *et al*.,^7^ we conclude that the diminished ionic strength could be due to neutralization of the ampholytes, i.e., of NH_4_^+^ ions into NH_3_, trimethylammonium into triethylamine, and OAc^−^ into HOAc. When the supercharged ions still follow the CRM, the alkali counter-ions condense on the nucleic acid, as depicted in molecular dynamics simulations of proteins in presence of Na^+^.^41^ The observation of more alkali adducts on higher charge states could mean that, in native supercharging by CRM, the higher charge states come from larger droplets, which upon evaporation reach a higher concentration of both the supercharging agent and of alkali counterions.

In 0.4% *m*-NBA, another supercharging mechanism can also take place, which involves the CEM and leads to even higher charge states. Because 0.4% volume *m*-NBA is above the solubility limit, phase separation inside the bulk solution combined with *m*-NBA affinity for nucleic acids can cause the formation of *m*-NBA aggregates that entrain the DNA. If such a charged aggregated is emitted by a parent droplet, it can produce denatured DNA structures at high charge state. Another (not mutually exclusive) scenario is ampholyte neutralization: if too many OAc^−^ ions are neutralized into HOAc in presence of high amounts of *m*-NBA, the DNA polyanions have to become the charge carriers on the droplet surface, and are thus more likely to take a CEM pathway.^6^

In summary, although we didn’t elucidate the supercharging mechanisms of nucleic acids in this single study, we presented few leads worth pursuing. We also hope that, by considering not only proteins, which usually carry both types of charge carriers, but also nucleic acids as representatives of polyanions in the negative mode, one can reach a more unified understanding of electrospray charging and supercharging.

## Acknowledgements

This work was funded by the Agence National de la Recherche (project ANR-18-CE29-0013 POLYnESI).

## Supporting information

Table of tuning parameters, melting curves, additional ESI-MS spectra and CCS distributions as described in the text (PDF).

## References

(1) Leney, A. C.; Heck, A. J. R. Native Mass Spectrometry: What Is in the Name? J. Am. Soc. Mass Spectrom. 2017, 28 (1), 5–13.

(2) Iribarne, J. V. On the Evaporation of Small Ions from Charged Droplets. J. Chem. Phys. 1976, 64 (6), 2287–2294.

(3) Fernandez de la Mora, J. Electrospray Ionization of Large Multiply Charged Species Proceeds via Dole’s Charged Residue Mechanism. Analytica Chimica Acta 2000, 406 (1), 93–104.

(4) Konermann, L.; Ahadi, E.; Rodriguez, A. D.; Vahidi, S. Unraveling the Mechanism of Electrospray Ionization. Anal. Chem. 2013, 85 (1), 2–9.

(5) Grandori, R. Origin of the Conformation Dependence of Protein Charge-State Distributions in Electrospray Ionization Mass Spectrometry. J. Mass Spectrom. 2003, 38 (1), 11–15.

(6) Khristenko, N.; Amato, J.; Livet, S.; Pagano, B.; Randazzo, A.; Gabelica, V. Native Ion Mobility Mass Spectrometry: When Gas-Phase Ion Structures Depend on the Electrospray Charging Process. J. Am. Soc. Mass Spectrom. 2019, 30 (6), 1069–1081.

(7) Loo, R. R. O.; Lakshmanan, R.; Loo, J. A. What Protein Charging (and Supercharging) Reveal about the Mechanism of Electrospray Ionization. J Am Soc Mass Spectrom 2014, 25 (10), 1675–1693.

(8) Iavarone, A. T.; Williams, E. R. Mechanism of Charging and Supercharging Molecules in Electrospray Ionization. J. Am. Chem. Soc. 2003, 125 (8), 2319–2327.

(9) Foley, E. D. B.; Zenaidee, M. A.; Tabor, R. F.; Ho, J.; Beves, J. E.; Donald, W. A. On the Mechanism of Protein Supercharging in Electrospray Ionisation Mass Spectrometry: Effects on Charging of Additives with Short- and Long-Chain Alkyl Constituents with Carbonate and Sulphite Terminal Groups. Analytica Chimica Acta: X 2019, 1, 100004.

(10) Iavarone, A. T.; Jurchen, J. C.; Williams, E. R. Supercharged Protein and Peptide Ions Formed by Electrospray Ionization. Anal. Chem. 2001, 73 (7), 1455–1460.

(11) Sterling, H. J.; Daly, M. P.; Feld, G. K.; Thoren, K. L.; Kintzer, A. F.; Krantz, B. A.; Williams, E. R. Effects of Supercharging Reagents on Noncovalent Complex Structure in Electrospray Ionization from Aqueous Solutions. J. Am. Soc. Mass Spectrom. 2010, 21 (10), 1762–1774.

(12) Sterling, H. J.; Williams, E. R. Origin of Supercharging in Electrospray Ionization of Noncovalent Complexes from Aqueous Solution. J. Am. Soc. Mass Spectrom. 2009, 20 (10), 1933–1943.

(13) Konermann, L.; Metwally, H.; Duez, Q.; Peters, I. Charging and Supercharging of Proteins for Mass Spectrometry: Recent Insights into the Mechanisms of Electrospray Ionization. Analyst 2019, 144 (21), 6157–6171.

(14) Xu, N.; Chingin, K.; Chen, H. Ionic Strength of Electrospray Droplets Affects Charging of DNA Oligonucleotides: ESI Charging of DNA Oligonucleotides. J. Mass Spectrom. 2014, 49 (1), 103–107.

(15) Brahim, B.; Alves, S.; Cole, R. B.; Tabet, J. C. Charge Enhancement of Single-Stranded DNA in Negative Electrospray Ionization Using the Supercharging Reagent Meta-Nitrobenzyl Alcohol. Journal of the American Society for Mass Spectrometry 2013, 24 (12), 1988–1996.

(16) Largy, E.; König, A.; Ghosh, A.; Ghosh, D.; Benabou, S.; Rosu, F.; Gabelica, V. Mass Spectrometry of Nucleic Acid Noncovalent Complexes. Chem. Rev. 2022, 122 (8), 7720–7839.

(17) Rhodes, D.; Lipps, H. J. G-Quadruplexes and Their Regulatory Roles in Biology. Nucleic Acids Res 2015, 43 (18), 8627–8637.

(18) Bowers, M. T.; Kemper, P. R.; von Helden, G.; van Koppen, P. A. M. Gas-Phase Ion Chromatography: Transition Metal State Selection and Carbon Cluster Formation. Science 1993, 260 (5113), 1446–1451.

(19) Clemmer, D. E.; Hudgins, R. R.; Jarrold, M. F. Naked Protein Conformations: Cytochrome c in the Gas Phase. J. Am. Chem. Soc. 1995, 117 (40), 10141–10142.

(20) Wyttenbach, T.; von Helden, G.; Bowers, M. T. Gas-Phase Conformation of Biological Molecules: Bradykinin. J. Am. Chem. Soc. 1996, 118 (35), 8355–8364.

(21) Cantor, C. R.; Warshaw, M. M.; Shapiro, H. Oligonucleotide Interactions. III. Circular Dichroism Studies of the Conformation of Deoxyoligonucleolides. Biopolymers 1970, 9 (9), 1059–1077.

(22) Mergny, J.-L.; Lacroix, L. Analysis of Thermal Melting Curves. Oligonucleotides 2003, 13 (6), 515–537.

(23) D’Atri, V.; Porrini, M.; Rosu, F.; Gabelica, V. Linking Molecular Models with Ion Mobility Experiments. Illustration with a Rigid Nucleic Acid Structure: Ion Mobility Calculations and Experiments. J. Mass Spectrom. 2015, 50 (5), 711–726.

(24) Marchand, A.; Livet, S.; Rosu, F.; Gabelica, V. Drift Tube Ion Mobility: How to Reconstruct Collision Cross Section Distributions from Arrival Time Distributions? Anal. Chem. 2017, 89 (23), 12674–12681.

(25) Do, N. Q.; Phan, A. T. Monomer–Dimer Equilibrium for the 5′–5′ Stacking of Propeller-Type Parallel-Stranded G-Quadruplexes: NMR Structural Study. Chemistry – A European Journal 2012, 18 (46), 14752–14759.

(26) Stewart, J. J. P. Optimization of Parameters for Semiempirical Methods VI: More Modifications to the NDDO Approximations and Re-Optimization of Parameters. J Mol Model 2013, 19 (1), 1–32.

(27) Frisch, M. J.; Trucks, G. W.; Schlegel, H. B.; Scuseria, G. E.; Robb, M. A.; Cheeseman, J. R.; Scalmani, G.; Barone, V.; Petersson, G. A.; Nakatsuji, H.; Li, X.; Caricato, M.; Marenich, A. V.; Bloino, J.; Janesko, B. G.; Gomperts, R.; Mennucci, B.; Hratchian, H. P.; Ortiz, J. V.; Izmaylov, A. F.; Sonnenberg, J. L.; Williams Ding, F.; Lipparini, F.; Egidi, F.; Goings, J.; Peng, B.; Petrone, A.; Henderson, T.; Ranasinghe, D.; Zakrzewski, V. G.; Gao, J.; Rega, N.; Zheng, G.; Liang, W.; Hada, M.; Ehara, M.; Toyota, K.; Fukuda, R.; Hasegawa, J.; Ishida, M.; Nakajima, T.; Honda, Y.; Kitao, O.; Nakai, H.; Vreven, T.; Throssell, K.; Montgomery Jr., J. A.; Peralta, J. E.; Ogliaro, F.; Bearpark, M. J.; Heyd, J. J.; Brothers, E. N.; Kudin, K. N.; Staroverov, V. N.; Keith, T. A.; Kobayashi, R.; Normand, J.; Raghavachari, K.; Rendell, A. P.; Burant, J. C.; Iyengar, S. S.; Tomasi, J.; Cossi, M.; Millam, J. M.; Klene, M.; Adamo, C.; Cammi, R.; Ochterski, J. W.; Martin, R. L.; Morokuma, K.; Farkas, O.; Foresman, J. B.; Fox, D. J. Gaussian 16 Rev. C.01, Wallingford, CT, 2016.

(28) Schlegel, H. B.; Millam, J. M.; Iyengar, S. S. Ab Initio Molecular Dynamics: Propagating the Density Matrix with Gaussian Orbitals. J. Chem. Phys. 2001, 114 (22), 7.

(29) Schlegel, H. B.; Iyengar, S. S.; Li, X. Ab Initio Molecular Dynamics: Propagating the Density Matrix with Gaussian Orbitals. III. Comparison with Born–Oppenheimer Dynamics. J. Chem. Phys. 2002, 117 (19), 12.

(30) Mesleh, M. F.; Hunter, J. M.; Shvartsburg, A. A.; Schatz, G. C.; Jarrold, M. F. Structural Information from Ion Mobility Measurements: Effects of the Long-Range Potential. J. Phys. Chem. 1996, 100 (40), 16082–16086

(31) Sterling, H. J.; Cassou, C. A.; Trnka, M. J.; Burlingame, A. L.; Krantz, B. A.; Williams, E. R. The Role of Conformational Flexibility on Protein Supercharging in Native Electrospray Ionization. Phys. Chem. Chem. Phys. 2011, 13 (41), 18288.

(32) Sterling, H. J.; Kintzer, A. F.; Feld, G. K.; Cassou, C. A.; Krantz, B. A.; Williams, E. R. Supercharging Protein Complexes from Aqueous Solution Disrupts Their Native Conformations. J. Am. Soc. Mass Spectrom. 2012, 23 (2), 191–200.

(33) Lomeli, S. H.; Yin, S.; Ogorzalek Loo, R. R.; Loo, J. A. Increasing Charge While Preserving Noncovalent Protein Complexes for ESI-MS. J. Am. Soc. Mass Spectrom. 2009, 20 (4), 593–596.

(34) Carter, D. V.; Charlton, P. T.; Fenton, A. H.; Housley, J. R.; Lessel, B. The Preparation and the Antibacterial and Antifungal Properties of Some Substituted Benzyl Alcohols. Journal of Pharmacy and Pharmacology 2011, 10 (Supplement_1), 149T-159T.

(35) Griffey, R. H.; Sasmor, H.; Greig, M. J. Oligonucleotide Charge States in Negative Ionization Electrospray-Mass Spectrometry Are a Function of Solution Ammonium Ion Concentration. J Am Soc Mass Spectrom 1997, 8 (2), 155–160.

(36) Gabelica, V.; Livet, S.; Rosu, F. Optimizing Native Ion Mobility Q-TOF in Helium and Nitrogen for Very Fragile No.ncovalent Structures. J. Am. Soc. Mass Spectrom. 2018, 29 (11), 2189–2198.

(37) Marchand, A.; Gabelica, V. Native Electrospray Mass Spectrometry of DNA G-Quadruplexes in Potassium Solution. J. Am. Soc. Mass Spectrom. 2014, 25 (7), 1146–1154.

(38) Miyoshi, D.; Karimata, H.; Sugimoto, N. Hydration Regulates Thermodynamics of G-Quadruplex Formation under Molecular Crowding Conditions. J. Am. Chem. Soc. 2006, 128 (24), 7957–7963.

(39) Miyoshi, D.; Sugimoto, N. Molecular Crowding Effects on Structure and Stability of DNA. Biochimie 2008, 90 (7), 1040–1051.

(40) Cassou, C. A.; Williams, E. R. Desalting Protein Ions in Native Mass Spectrometry Using Supercharging Reagents. Analyst 2014, 139 (19), 4810–4819.

(41) Metwally, H.; McAllister, R. G.; Popa, V.; Konermann, L. Mechanism of Protein Supercharging by Sulfolane and m -Nitrobenzyl Alcohol: Molecular Dynamics Simulations of the Electrospray Process. Anal. Chem. 2016, 88 (10), 5345–5354.

