## supporting information for "Negative Electrospray Supercharging Mechanisms of Oligonucleotide Single Strands and G-quadruplexes"

**Table of Contents**


| | Description | Page Number |
|---|---|---|
| **Figure S1** | Melting curves of ***20G*** | 2 |
| **Figure S2** | Melting curves of ***24G*** | 3 |
| **Figure S3** | Collision cross sections ($^{DT}CCS_{He}$) of the ***20G***$^{5-}$ (experimental and theoretical) | 4 |
| **Figure S4** | Comparison of CSD and CCSDs in 7-, 8-, and 9- charge states of ***20G*** | 5 |
| **Figure S5** | CSDs of ***24G*** and ***24nonG4*** in different SCAs | 6 |
| **Figure S6** | CCSDs of ***24G***(A) and ***24nonG4***(B) in different SCAs | 7 |
| **Table S1** | Tuning parameters for COMP and OPT conditions in the Post-IMS Region | 8 |
| **Figure S7** | Formation of *m*-NBA adducts with ***TG4T*** in extra soft post-IMS conditions with 0.1% of *m*-NBA in 100 mM aqueous $NH_4OAc$ | 9 |
| **Figure S8** | CSD of ***20nonG*** in (A) 100 mM TMAA (no $K^+$), (B) 100 mM $NH_4OAc$ (no $K^+$) | 10 |
| **Figure S9** | Filtering of ESI-MS data thanks to ion mobility, based on the charge states | 11 |
| **Figure S10** | CCSD of ***20G*** in 100 mM TMAA and 0.3 mM $K^+$ | 12 |
| **Figure S11** | Non-specific $K^+$ adducts of ***20nonG*** with SCA in 100 mM $NH_4OAc$ and 0.3 mM $K^+$ | 13 |
| **Figure S12** | Non-specific $K^+$ adducts ***20nonG*** with SCA and 0.3 mM $K^+$ in 100 mM and 1 mM TMAA | 14 |

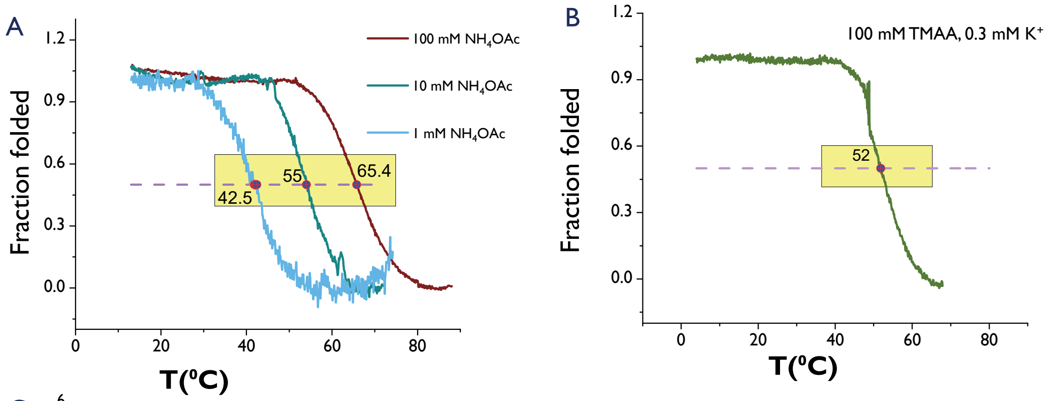


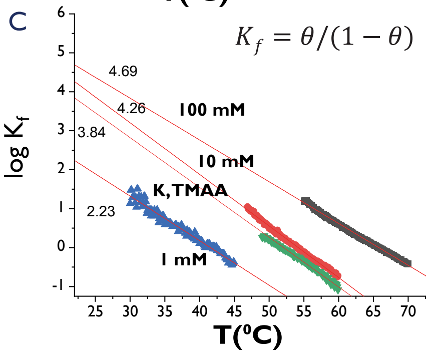


D

| % Folded (22°C) | Ionic strength |
|---|---|
| 99.99% | 100 mM NH4OAc |
| 99.98% | 100 mM TMAA, 0.3 mM K$^+$ |
| 99.99% | 10 mM NH4OAc |
| 99.41% | 1 mM NH4OAc |

**Figure S1.** Melting curve of ***20G*** showing that the structures are folded in solution at room temperature. A) Results in 100, 10, and 1 mM aqueous $NH_4OAc$ solution. B) Result in 100 mM aqueous TMAA and 0.3 mM KOAc. C) Plot of log $K_f$ as a function of the temperature. D) % folded at room temperature.

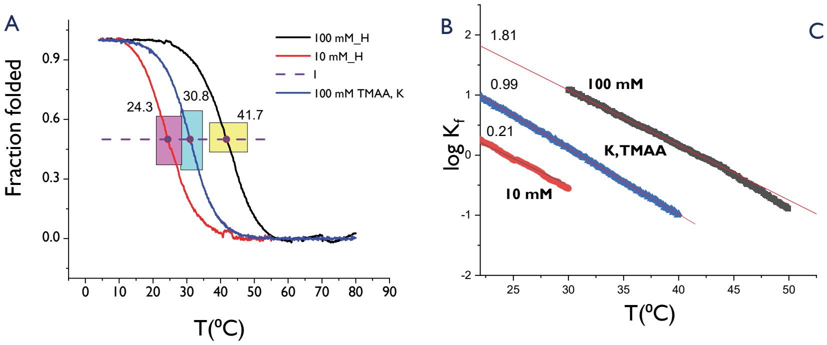


C

| % Folded (22°C) | Ionic strength |
|---|---|
| 98.48% | 100 mM NH4OAc |
| 90.90% | 100 mM TMAA, 0.3 mM $K^+$ |
| 66.66% | 10 mM NH4OAc |

**Figure S2.** Melting curves of ***24G*** showing that the structures are folded in solution at room temperature. A) Results in 100, 10 aqueous $NH_4OAc$, and 100 mM aqueous TMAA, 0.3 mM KOAc. B) Plot of log $K_f$ as a function of the temperature. C) % folded at room temperature.

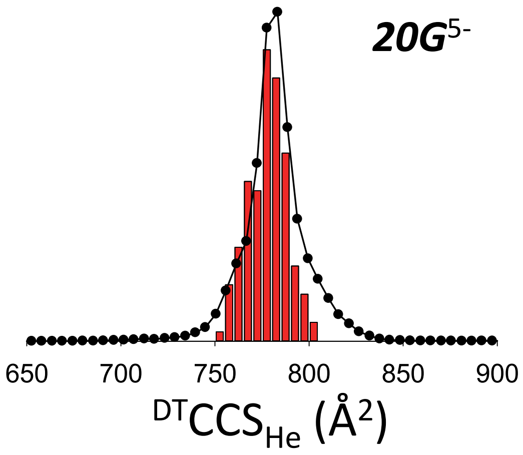


**Figure S3.** Dots: collision cross section distribution ($^{DT}CCS_{He}$) of the ***20G***$^{5-}$ measured by drift tube ion mobility MS. The corresponding theoretical CCSs calculated from snapshots extracted from molecular dynamics simulations is displayed as histograms. Semi-empirical level (PM7) is used for the MD.

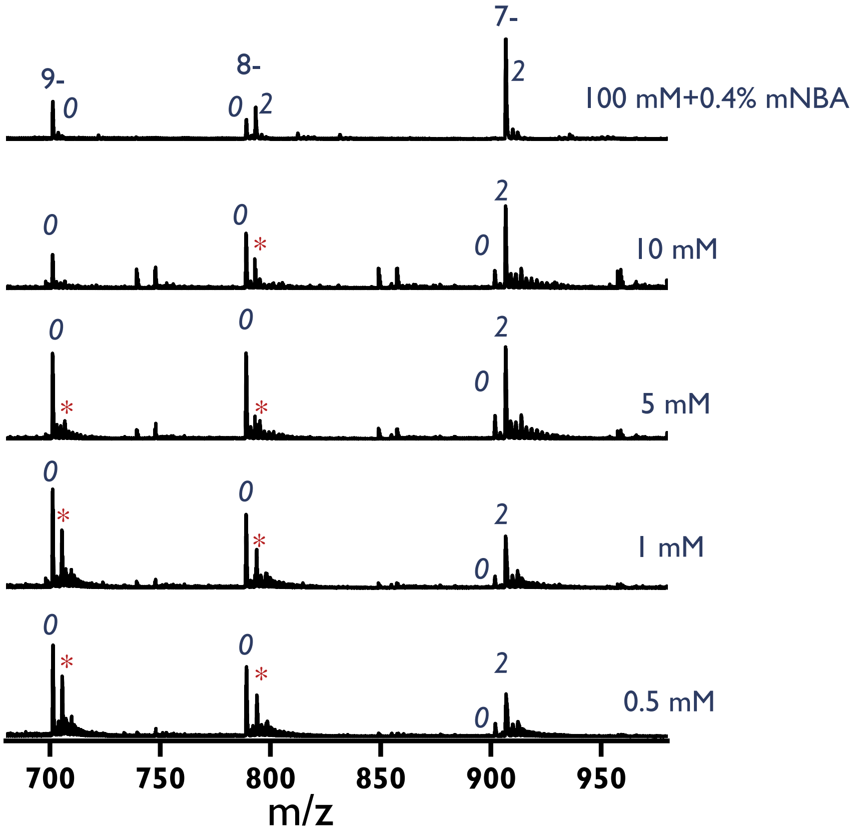


**Figure S4.** ESI-MS spectra (7-, 8-, and 9- charge states) for ***20G*** with 0.4% *m*-NBA added to 100 mM $NH_4OAc$, compared to spectra acquired at lower [$NH_4OAc$]. The cation loss starts at charge state 8- in presence of *m*-NBA in 100 mM aqueous $NH_4OAC$ solution, and from 7- for solutions prepared at lower ionic strength in absence of *m*-NBA. *0* indicates no specific $NH_4^+$ bound and *2* indicates 2 specific $NH_4^+$ ions bound. * indicates a $K^+$ adduct, issued from unwanted contamination.

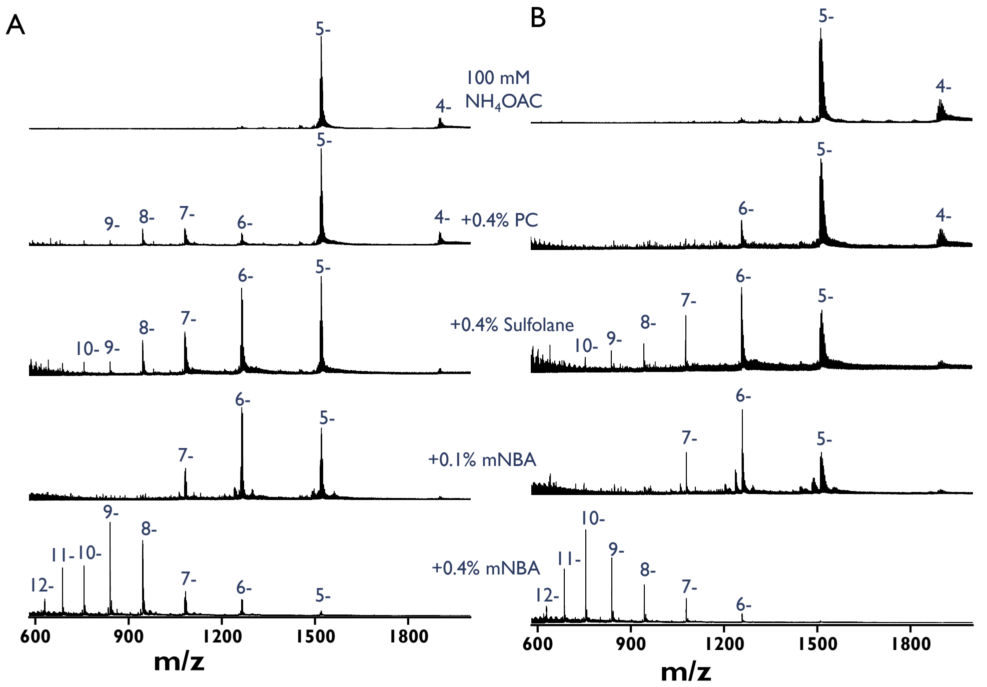


**Figure S5.** ESI-MS spectra of ***24G*** (A) and ***24nonG4*** (B) without and with different SCAs (0.4% PC, 0.4% sulfolane, 0.1% *m*-NBA and 0.4% *m*-NBA) in 100 mM aqueous $NH_4OAc$.

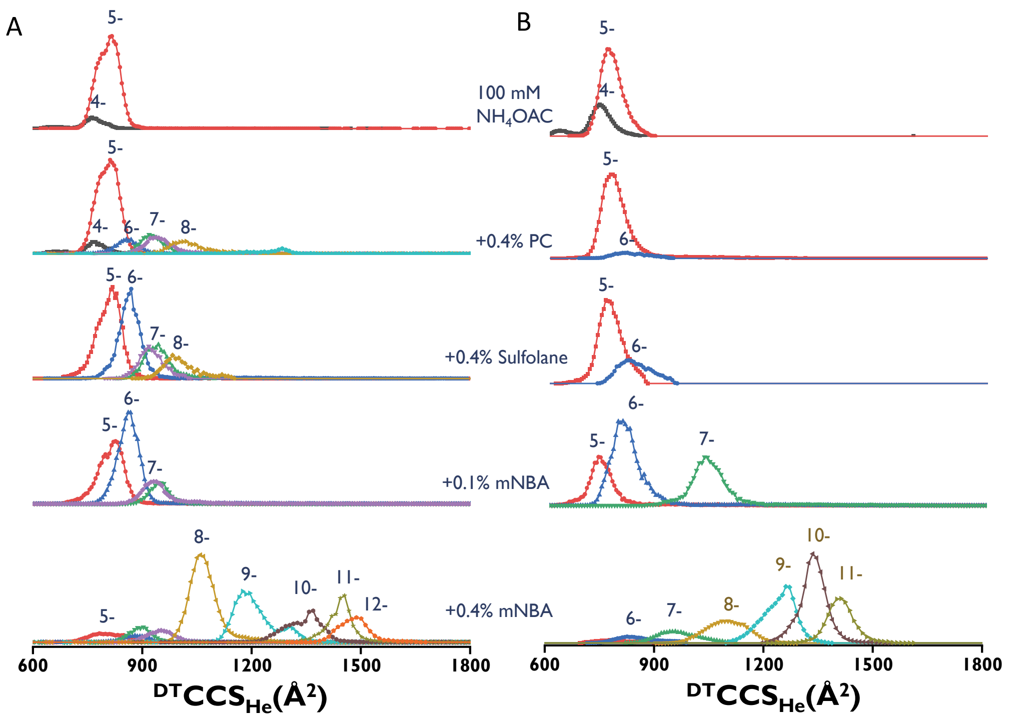


**Figure S6.** Collision cross section distributions of ***24G*** (A) and ***24nonG4*** (B) without and with different SCAs (0.4% PC, 0.4% sulfolane, 0.1% *m*-NBA and 0.4% *m*-NBA) in 100 mM aqueous $NH_4OAc$.

**Table S1.** Tuning parameters for soft and extra-soft post-IMS conditions in negative mode. These settings correspond to the COMP and OPT tuning conditions described in: Gabelica, V.; Livet, S.; Rosu, F. Optimizing Native Ion Mobility Q-TOF in Helium and Nitrogen for Very Fragile Noncovalent Structures. *J. Am. Soc. Mass Spectrom.* **2018**, *29* (11), 2189–2198. https://doi.org/10.1007/s13361-018-2029-4.

| **Parameter** | **Soft post-IMS** | **Extra soft post-IMS** |
|---|---|---|
| IM rear funnel: rear funnel exit | -35 V | -26 V |
| IM rear funnel: IM Hex Entrance | -32 V | -24 V |
| IM rear funnel: IM Hex Delta | -3 V | -2 V |
| Optics 1: Oct Entrance Lens | -27 V | -21 V |
| Optics 1: Oct 1 DC | -25 V | -20 V |
| Optics 1: Lens 1 | -23 V | -19 V |
| Quad: Quad DC | − 21 V | − 18 V |
| Quad: postfilter DC | − 21 V | − 17 V |
| Cell: gas flow | 20 psi | 20 psi |
| Cell: cell entrance | − 20 V | − 16 V |
| Cell: Hex DC | − 20 V | − 16 V |
| Cell: Hex Delta | 3 V | 3 V |
| Cell: Hex2 DC | − 14.6 V | − 12 V |
| Cell: Hex2 DV | 1.5 V | 1 V |
| Optics 2: Hex3 DC | − 12.9 V | − 11 V |
| Extractor: ion focus | − 10 V | − 10 V |

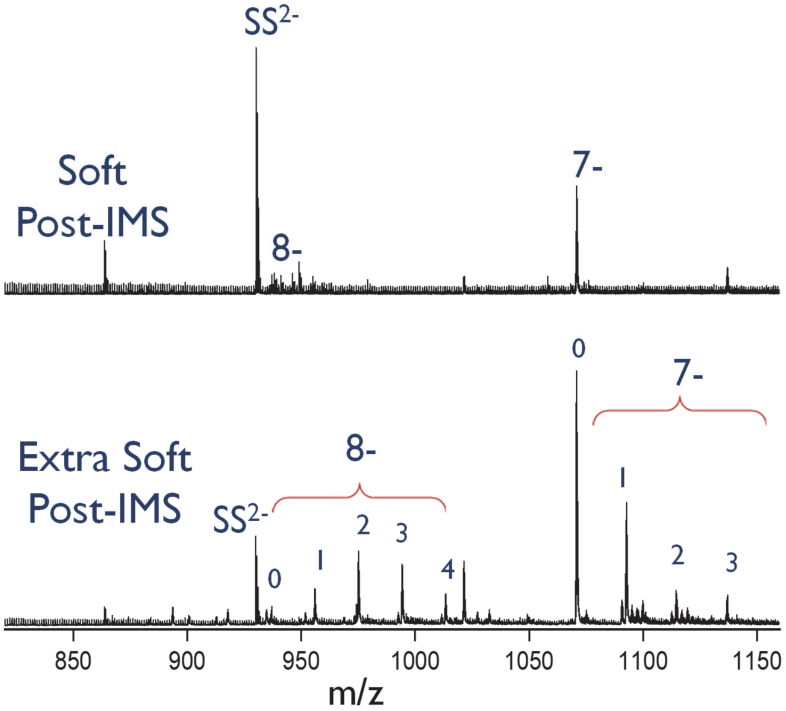


**Figure S7.** Formation of *m*-NBA adducts (number of adducts indicated in the figure) with the G-quadruplex of ***TG4T*** detected on charge states 8- and 7- in extra soft post-IMS conditions. Experiments were done using 0.1% of *m*-NBA in 100 mM aqueous $NH_4OAc$. The changes made in the tuning for post-IMS are listed in Table S1.

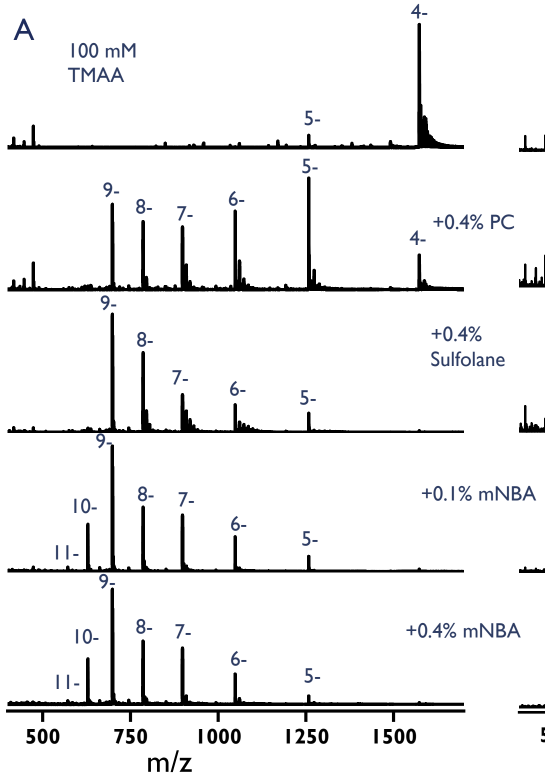


**Figure S8.** ESI-MS spectra of ***20nonG*** in (A) 100 mM TMAA (no $K^+$) and (B) 100 mM $NH_4OAc$ (no $K^+$).

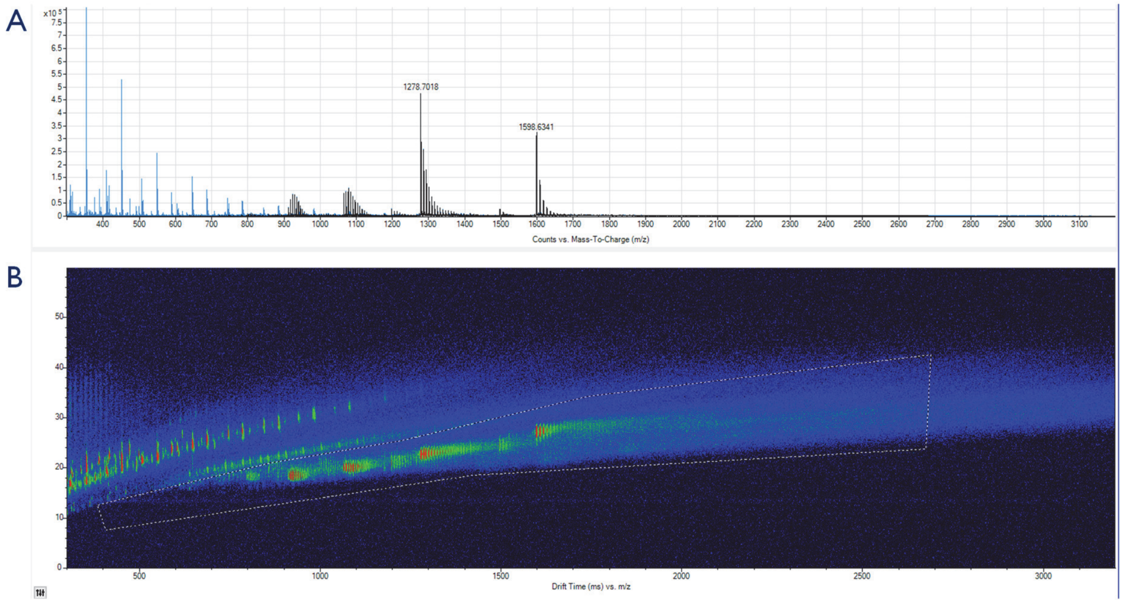


**Figure S9.** Extraction of MS data in IM-MS browser. A) ESI-MS spectrum of ***20G*** in 100 mM TMAA, 0.3 mM $K^+$: blue+black = full specrum; black = filtered spectrum. B) Drift time vs. *m/z* plot of ***20G*** in 100 mM TMAA, 0.3 mM $K^+$. White dotted line is drawn (B, 2D plot) to exclude the presence of single and doubly charged background from the MS data. The resultant MS data of highly charged oligonucleotide ions is shown by black spectrum and original raw data is shown by blue spectrum in A. This is an example for results in 0.4% PC. Other results in Figure 6 were obtained with a similar procedure.

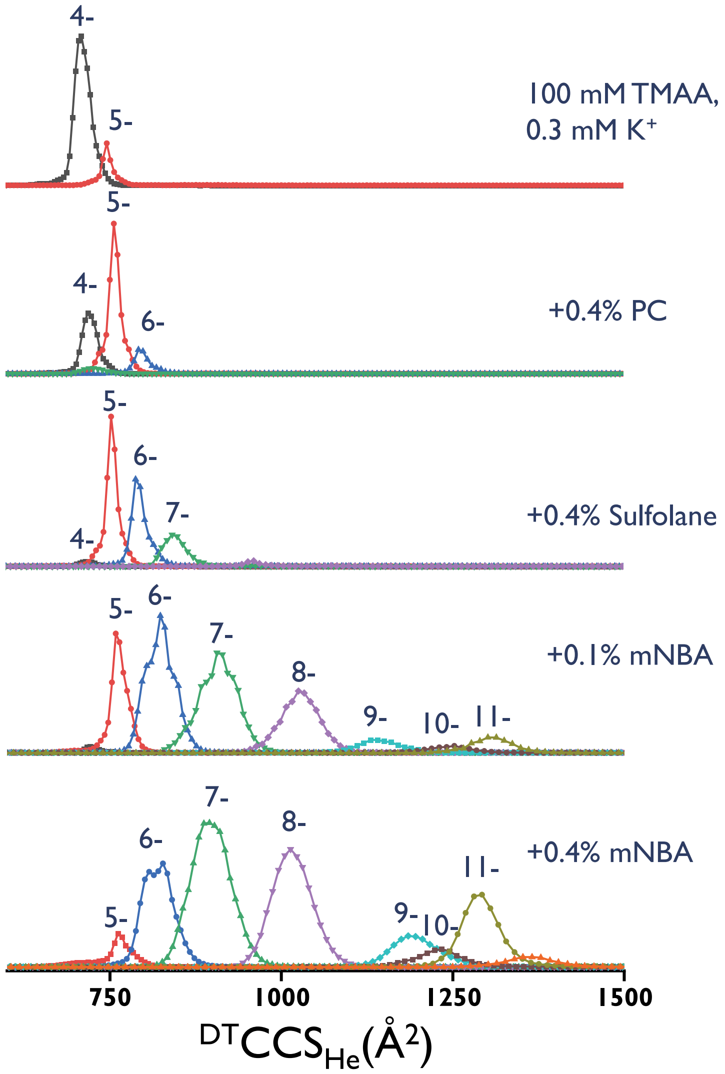


**Figure S10.** Collision cross section distributions of ***20G*** in 100 mM TMAA and 0.3 mM KOAc.

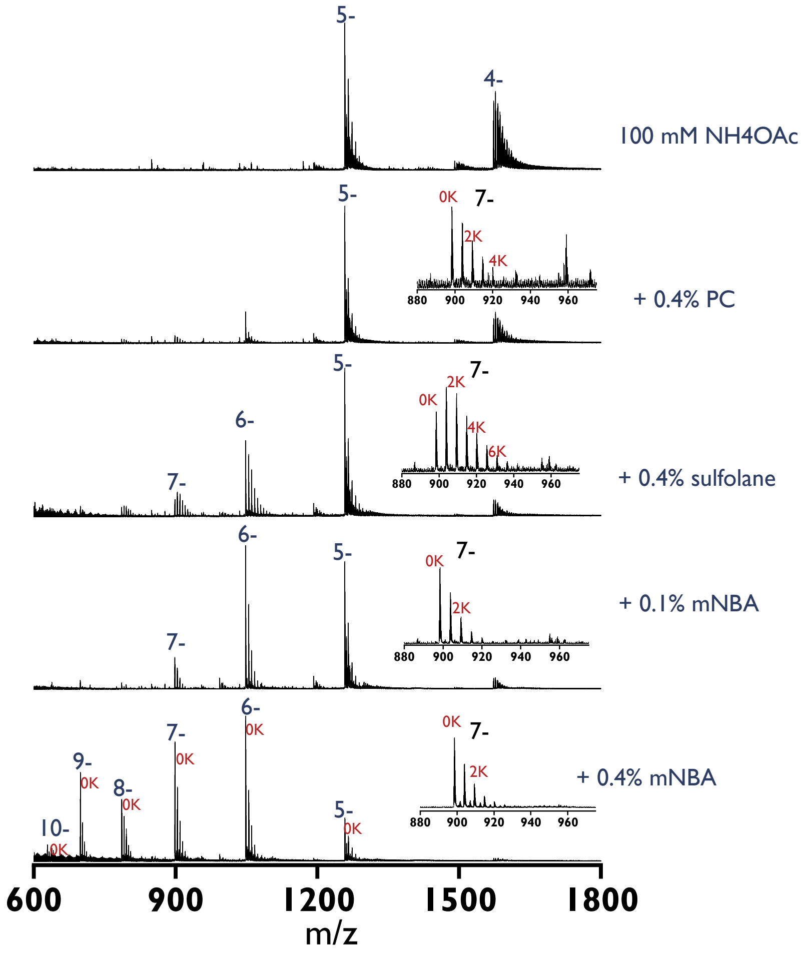


**Figure S11.** Formation of non-specific $K^+$ adducts of ***20nonG*** with SCA in 100 mM $NH_4OAc$ and 0.3 mM $K^+$.

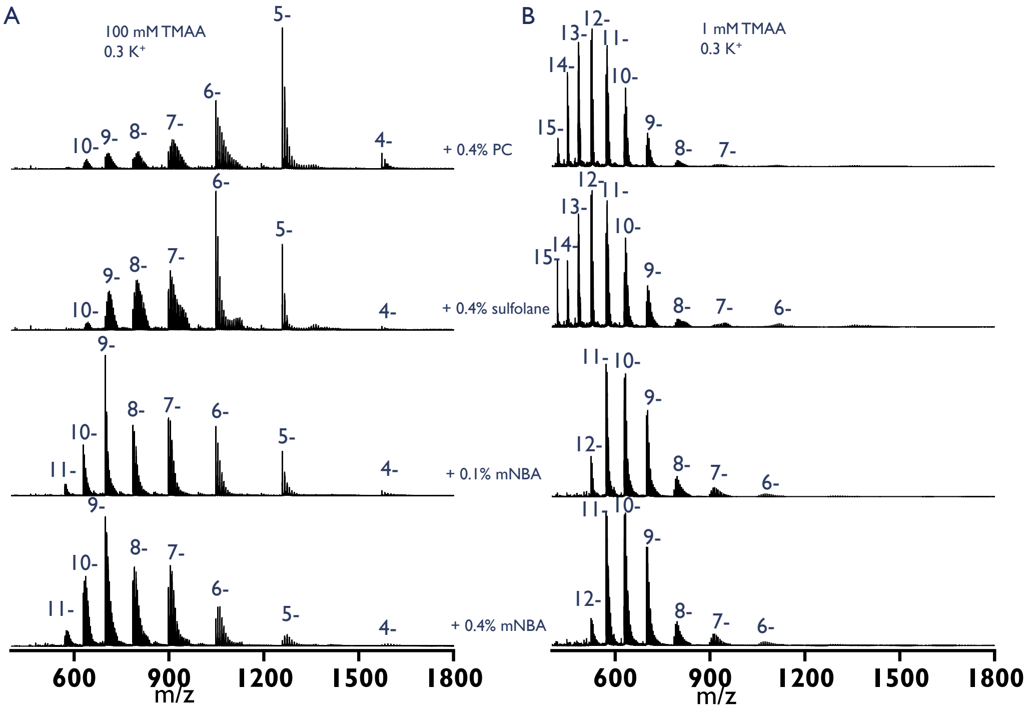


**Figure S12.** ESI-MS spectra showing the formation of non-specific $K^+$ adducts of ***20nonG*** with SCA and 0.3 mM KOAc, 100 mM TMAA (left panel) and 1 mM TMAA (right panel).
